# Non-micronized and micronized curcumin do not prevent the behavioral and neurochemical effects induced by acute stress in zebrafish

**DOI:** 10.1101/2021.10.11.463974

**Authors:** Adrieli Sachett, Matheus Gallas-Lopes, Radharani Benvenutti, Matheus Marcon, Amanda M. Linazzi, Gean P. S. Aguiar, Ana P. Herrmann, J. Vladimir Oliveira, Anna M. Siebel, Angelo Piato

## Abstract

Curcumin, a polyphenol extracted from the rhizome of *Curcuma longa* L. (Zingiberaceae), presents neuroprotective properties and can modulate neuronal pathways related to mental disorders. However, curcumin has low bioavailability, which can compromise its use. The micronization process can reduce the mean particle diameter and improve this compound’s bioavailability and therapeutic potential. In this study, we compared the behavioral (in the open tank test, OTT) and neurochemical (thiobarbituric acid reactive substances (TBARS) and non-protein thiols (NPSH) levels) effects of non-micronized curcumin (CUR, 10 mg/kg, i.p.) and micronized curcumin (MC, 10 mg/kg, i.p.) in adult zebrafish subjected to 90-minute acute restraint stress (ARS). ARS increased the time spent in the central area and the number of crossings and decreased the immobility time of the animals. These results suggest an increase in locomotor activity and a decrease in thigmotaxis behavior in the OTT. Furthermore, ARS also induced oxidative damage by increasing TBARS and decreasing NPSH levels. ARS-induced behavioral and biochemical effects were not blocked by any curcumin preparation. Therefore, we suppose that curcumin does not have anti-stress effects on the ARS in zebrafish.

## Introduction

The term stress refers to the set of responses triggered by the body in a situation assumed as threatening, which occurs due to a stressor. These responses are triggered by activation of the autonomic nervous system and hypothalamic-pituitary-adrenocortical (HPA) axis, which results in the release of catecholamines and glucocorticoids. It aims to adapt the system to a given demand, adapt and return to the basal level of internal balance, called homeostasis. The persistence of these responses, even when the stress stimulus ends, indicates a failed adaptation, which can affect an individual’s health status and result in mental disorders such as anxiety and depression [1–5]. Stress can lead to the depletion of the body’s adaptive responses through complex neurobiological changes involving oxidative stress, neuroinflammation, neurotransmitters signaling disfunction, and excitotoxicity that can ultimately affect neurocircuits that regulate behavior, especially behaviors relevant to decreased motivation (anhedonia), avoidance, and alarm (anxiety), which characterize several neuropsychiatric disorders [6]. Although effective treatments for these conditions are available, a significant number of patients do not adequately respond to the therapy, further contributing to the global burden of these mental diseases. Therefore, it is essential to study innovative treatments that modulate targets related to these psychopathologies [6, 7].

Zebrafish stress response is mediated by the hypothalamic-pituitary-interrenal (HPI) axis that is functionally and structurally homologous to the mammalian HPA axis [8]. In response to an acute stressor (for example, restraint, alarm pheromone, net chasing), zebrafish exhibit a complex behavioral and physiological repertoire including anxiety, locomotor disturbance, increase in thigmotaxis behavior, cognitive impairment, hypercortisolemia, and oxidative status imbalance [9–21]. These effects were blocked by antidepressants/ anxiolytics [9, 16], antipsychotics [17], and other compounds [18, 19, 21].

Curcumin, a compound extracted from the roots of the ground turmeric (*Curcuma longa* L. Zingiberaceae), presented antioxidant, anti-inflammatory, neuroprotective, immunomodulatory, anxiolytic, and antidepressant effects in several pre-clinical and clinical studies [7, 22–26]. However, curcumin has low bioavailability, poor absorption, rapid metabolism, and quick systemic elimination, which compromise its therapeutic use for neuropsychiatric disorders [27, 28]. Supercritical fluid micronization technology (SEDS) is an approach that has been using to modify material properties by reducing particle size, increasing dissolution rate and solubility, as well as modifying the crystal structure of the compound when compared to non-micronized compounds [29, 30]. These changes can potentially increase bioavailability. Therefore, this study aimed to compare the effects of non-micronized curcumin and micronized curcumin on the behavioral and biochemical parameters in adult zebrafish submitted to acute stress.

## Materials and methods

### Drugs

Curcumin was obtained from Sigma-Aldrich® (CAS 458-37-7) (St. Louis, MO, USA). Curcumin micronization was carried out at the Laboratory of Thermodynamics and Supercritical Technology (LATESC) of the Department of Chemical and Food Engineering (EQA) at Universidade Federal de Santa Catarina (UFSC), with the solution enhanced dispersion by supercritical fluids (SEDS) according to [31]. Both curcumin preparations were dissolved in 1% DMSO (dimethyl sulfoxide anhydrous) obtained from Sigma-Aldrich® (CAS 67-68-5) and diluted in injection water (Samtec Biotecnologia®, SP, Brazil) acquired from a commercial supplier. Reagents used for biochemical assays were obtained from Sigma Aldrich (St. Louis, MO, USA), 5,5’-dithiobis (2-nitrobenzoic acid) (CAS Number 69-78-3), thiobarbituric acid (CAS Number: 504-17-6), and trichloroacetic acid (CAS Number: 76-03-9). Absolute ethanol (CAS Number: 64-17-5) was obtained from Merck KGaA (Darmstadt, Germany).

### Animals

All procedures were approved by the institutional animal welfare and ethical review committee at the Universidade Federal do Rio Grande do Sul (UFRGS) (approval #35279/2018). The animal experiments are reported in compliance with the ARRIVE guidelines 2.0 [32]. Experiments were performed at the Laboratory of Psychopharmacology and Behavior (LAPCOM) of the Department of Pharmacology at UFRGS, using 144 male and female (50:50 ratio) short-fin wild-type zebrafish, 6 months old, weighing 300 to 400 mg. Adult animals were obtained from the colony established in the Biochemistry Department at UFRGS and maintained for at least 15 days before tests in our animal facility (Altamar, SP, Brazil) in 16-L home tanks (40 × 20 × 24 cm) with non-chlorinated water kept under constant mechanical, biological, and chemical filtration at a maximum density of two animals per liter. Tank water fulfilled the controlled conditions required for the species (26 ± 2 °C; pH 7.0 ± 0.3; dissolved oxygen at 7.0 ± 0.4 mg/L; total ammonia at <0.01 mg/L; total hardness at 5.8 mg/L; alkalinity at 22 mg/L CaCO_3_; and conductivity of 1500–1600 μS/cm). The animals were maintained in a light/dark cycle of 14/10 hours and food was provided twice a day (commercial flake food (Poytara®, Brazil) plus the brine shrimp *Artemia salina*). After the tests, animals were euthanized by hypothermic shock according to the AVMA Guidelines for the Euthanasia of Animals [33]. Briefly, animals were exposed to chilled water at a temperature between 2 and 4 °C for at least 2 min after loss of orientation and cessation of opercular movements, followed by decapitation as a second step to ensure death.

### Drug administration

Intraperitoneal (i.p.) injections were applied using a Hamilton Microliter™ Syringe (701N 10 μL SYR 26s/2“/2) × Epidural catheter 0.45 × 0.85 mm (Perifix®-Katheter, Braun, Germany) × Gingival Needle 30G/0.3 × 21 mm (GN injecta, SP, Brazil). The animals’ weight was checked 24 hours before the treatment and an average between the weights of each tank was used to calculate the injection volume (1 μL/100 mg of animal weight). The animals were anesthetized by immersion in a solution of tricaine (300 mg/L, CAS number 886-86-2) until postural loss and reduction of respiratory rate. The anesthetized fish were gently placed in a sponge soaked in water placed inside a petri dish, with the abdomen facing up and the fishs’ head positioned on the sponges’ hinge. The needle was inserted parallel to the spine in the abdomens’ midline posterior to the pectoral fins. This procedure was conducted in approximately 10 seconds. The behavioral tests took place 90 minutes after the injection.

### Experimental design

The experimental design is presented in Fig. 1. Initially, animals were treated with 1% DMSO, 10 mg/kg CUR, or 10 mg/kg MC (n= 24).These doses were standardized in our previous study [31]. After the treatment, the experimental groups were subdivided into control (non-stressed group, S-) or acute restraint stress (ARS) (stressed group, S+). The stress protocol was conducted as described previously [10, 12, 15, 20, 21]. Fish in the stressed group were gently placed into 1.8-mL cryogenic tubes (Corning®) with openings at both ends (to allow adequate water circulation) in a 16-L tank for 90 min. Non-stressed groups were transferred to an identical 16-L tank for 90 min. After the acute stress protocol, the animals were individually transferred to the OTT, and behavioral parameters were quantified. Immediately after the OTT, fish were euthanized, and the brain was dissected and homogenized for the biochemical assays. The animals were allocated to the experimental groups following block randomization procedures to counterbalance the sex, the two different home tanks, and the test arenas between the groups. Animal behavior was video recorded and analyzed with the ANY-Maze tracking software (Stoelting Co., Wood Dale, IL, USA) by researchers blinded to the experimental groups. All tests were performed between 08:00 and 12:00 a.m. The sex of the animals was confirmed after euthanasia by dissecting and analyzing the gonads.

**Fig 1.**
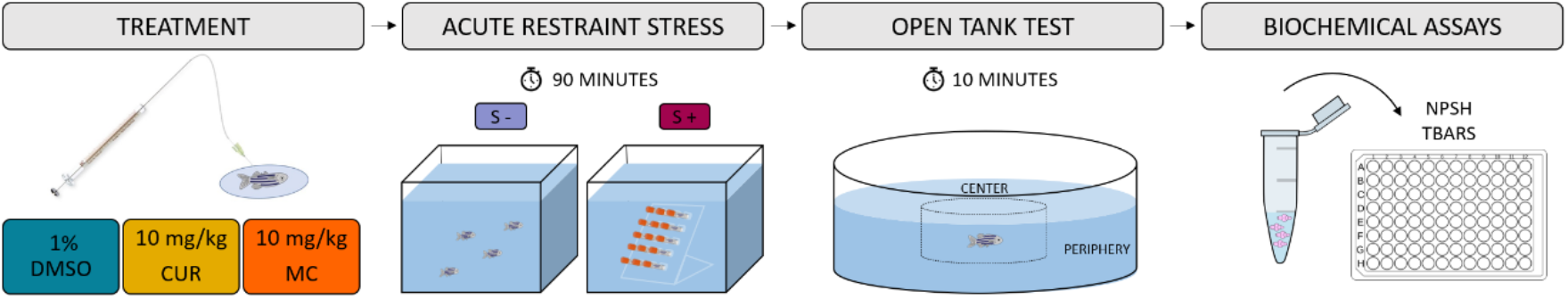
Experimental design. The treatments 1% DMSO, 10 mg/kg CUR, or 10 mg/kg MC were injected intraperitoneally. Afterward, zebrafish were subjected to acute restraint stress for 90 minutes. Zebrafish from the control group remained in an identical tank and were not submitted to stress. Subsequently, the animals were tested in the open tank test. Immediately after the behavioral test, each animal was euthanized and the brain dissected to performed the biochemical assays. DMSO (dimethyl sulfoxide), CUR (curcumin), MC (micronized curcumin), NPSH (non-protein thiols) and TBARS (thiobarbituric acid reactive substances)

### Open tank test (OTT)

The OTT was conducted as described previously [10, 31, 34]. Animals were individually placed in the center of a circular arena made of opaque white plastic (24 cm diameter, 8 cm walls, 2 cm water level) and recorded for 10 min. The apparatus was virtually divided into two areas for video analysis: the central (12 cm in diameter) and the periphery areas. Videos were recorded from the top view. The following parameters were quantified: total distance traveled (m), number of crossings (transitions between the areas of the tank), absolute turn angle (°), immobility time, time spent, and entries in the center area of the tank.

### Neurochemical assays

For each independent sample, four brains were pooled (n=6) and homogenized in 600 μL of phosphate-buffered saline (PBS, pH 7.4, Sigma-Aldrich) and centrifuged at 10,000 g at 4 °C in a cooling centrifuge; the supernatants were collected and kept in microtubes on ice until the assays were performed. The detailed protocol for prepare brain tissue samples is available at protocols.io [35]. The protein content was quantified according to the Coomassie blue method using bovine serum albumin (Sigma-Aldrich) as a standard [36]. The detailed protocol for protein quantification is available at protocols.io [37].

### Non-protein thiols (NPSH)

The content of NPSH in the samples was determined by mixing equal volumes of the brain tissue preparation (50 μg of proteins) and trichloroacetic acid (TCA, 6%), centrifuging the mix (10,000 g, 10 min at 4 °C), the supernatants were added to TFK (1 M) and DTNB (10 mM) and the absorbance was measured at 412 nm after 1 h. The detailed protocol is available at protocols.io [38].

### Thiobarbituric acid reactive substances (TBARS)

The lipid peroxidation was evaluated by quantifying the production of TBARS. Samples (50 μg of proteins) were mixed with TBA (0.5%) and TCA (20%) (150 μL). The mixture was heated at 100 °C for 30 min. The absorbance of the samples was determined at 532 nm in a microplate reader. MDA (2 mM) was used as the standard. The detailed protocol is available at protocols.io [39].

### Statistical analysis

We calculated the sample size to detect an effect size of 0.35 for the interaction between stress and treatment with a power of 0.9 and an alpha of 0.05 using G*Power 3.1.9.7 for Windows. The total distance traveled was defined as the primary outcome. The total sample size was 144, n= 24 animals per experimental group.

The normality and homogeneity of variances were confirmed for all data sets using D’Agostino-Pearson and Levene tests, respectively. Results were analyzed by two-way ANOVA. The outliers were defined using the ROUT statistical test and were removed from the analyses. This resulted in 6 outliers being removed from the OTT test (1 animal from CUR S-, 2 animals from DMSO S+ and 3 animals from MC S+ groups). Moreover, one animal showed 0 distance in OTT from the MC S+ group and was removed from the test. The tank and sex effects were tested in all comparisons and no effect was observed, so the data were pooled.

Data are expressed as mean ± standard deviations of the mean (S.D.). The level of significance was set at p<0.05. Data were analyzed using IBM SPSS Statistics version 27 for Windows and the graphs were plotted using GraphPad Prism version 8.0.1 for Windows.

## Results

### Behavioral parameters

The effects of CUR and MC on behavioral parameters in zebrafish submitted to ARS in the open tank test (OTT) are presented in Fig 2. Two-way ANOVA revealed that acute stress increased the number of crossings (Fig. 2B) and decreased the time immobile (Fig. 2D). Also, ARS increased the time spent in the center area (Fig. 2E), indicating a decrease in the thigmotaxic behavior. Both CUR and MC did not prevent the effects of acute stress on these behavioral parameters. There was no statistical difference in the parameters of total distance traveled, absolute turn angle, and center entries.

**Fig. 2.**
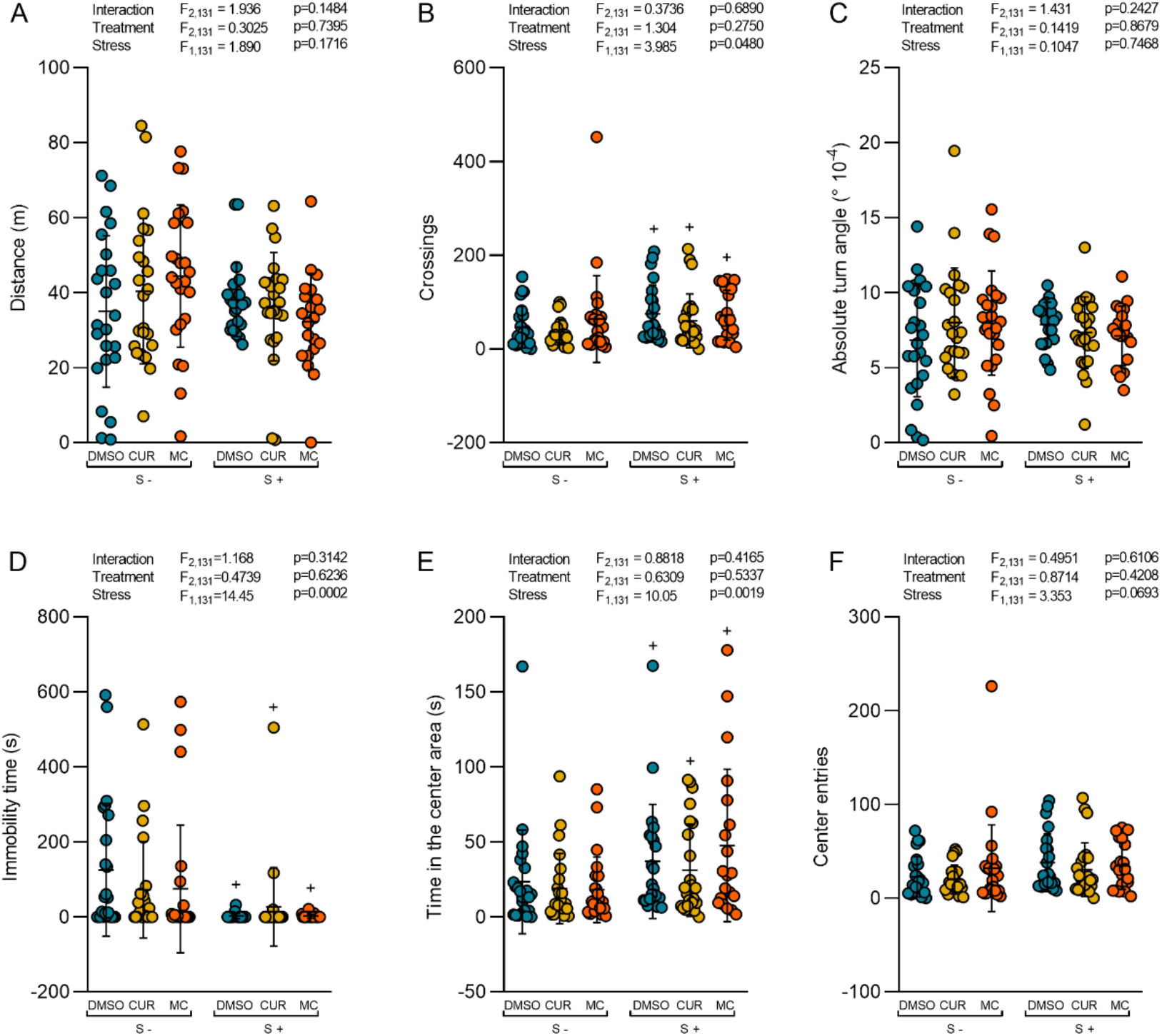
Effects of CUR and MC (10 mg/kg) on behavioral parameters in zebrafish submitted to the acute restraint stress. (A) total distance traveled, (B) number of crossings, (C) absolute turn angle, (D) immobility time, (E) time in the center area, and (F) center entries. Data are expressed as mean ± S.D. Two-way ANOVA. n=20-24. ^+^p<0.05 stress effect. DMSO (dimethyl sulfoxide), CUR (curcumin), MC (micronized curcumin)

### Neurochemical parameters

The effects of CUR and MC on oxidative status parameters in zebrafish brains submitted to ARS are presented in Fig 3. Two-way ANOVA revealed the main effect of stress on NPSH (Fig. 3A) and TBARS (Fig. 3B) levels, decreasing and increasing, respectively. These results indicate ARS provoked oxidative stress in the zebrafish brain. Both CUR and MC were not able to prevent these effects.

**Fig. 3.**
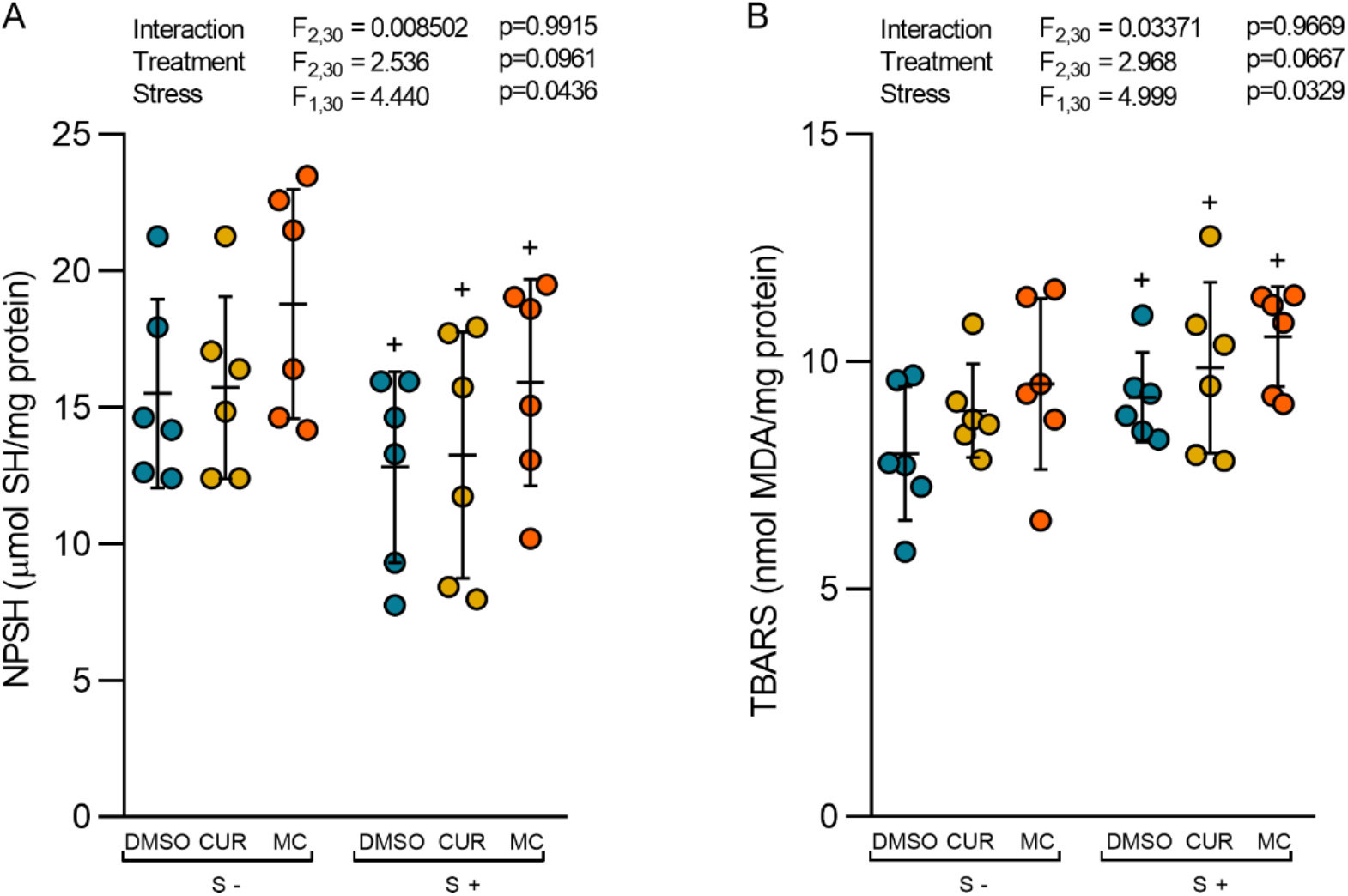
Effects of CUR and MC (10 mg/kg) on neurochemical parameters in zebrafish submitted to the acute restraint stress. (A) NPSH and (B) TBARS. Data are expressed as mean ± S.D. Two-way ANOVA. n=6. ^+^p<0.05 stress effect. DMSO (dimethyl sulfoxide), CUR (curcumin), MC (micronized curcumin), NPSH (non-protein thiols), TBARS (thiobarbituric acid reactive substances)

## Discussion

In this study, ARS increased locomotion and decreased thigmotaxis behavior in the open tank test, and induced oxidative damage in the zebrafish brain. Both curcumin preparations were not able to prevent the effects of ARS on the behavioral and neurochemical parameters tested.

The acute restraint stress model merges both emotional and physical aspects of stress and is widely used to study the impact of stress on disease processes and in stress-associated pathological conditions [40]. In response to an acute stressor, zebrafish may display fear/anxiety-like behaviors, such as altered locomotion (e.g., increased distance and average speed spent in the tank) as well as memory deficits and increase of cortisol levels [10, 11, 13, 16, 20, 21].

Traditionally, an increase in the time and exploration of aversive zones (bolder behavior) in open field tests are usually regarded as anxiolytic effects [41, 42]. Acute restraint stress has been reported to alter locomotor activity and cause anxiety-like behaviors in zebrafish [10, 20, 21]. In this study, ARS increased locomotion and decreased thigmotaxis in the open tank test. These findings, therefore, could suggest that acute stress may momentarily relieve anxiety in zebrafish, which appears counterintuitive but is not unprecedented in the literature [11]. The same was observed in zebrafish submitted to the acute restraint stress (15 minutes), which showed an increase in the number of the crossings, and spent significantly less time in the external area (periphery) than non-stressed controls in the OTT [11]. Also, acute stress has been previously shown to affect behavioral patterns in rodents, in a nonconventional manner as well. For instance, in a study in rodents, acute immobilization (120 minutes) produced anxiety-like behavior in the elevated plus-maze, which is reflected by hyperactivity in the open field test [40]. Immediate exposure to stress has been reported to induce anxiogenic behavior that may result in an excitable and irritable state leading to impaired performance [40, 43]. Furthermore, acute stress paradigms including restraint stress can cause a reduction in anxiety-like behaviors such as risk assessment behaviors (head poking and stretch attempts) and at the same time cause increases in the frequency of direct entries in the inner/anxiogenic zone of a novel environment such as the open field [11]. The reason underlying this phenomenon is unclear. One possibility is that acute stress may momentarily alter cognitive function, which may, in turn, affect the selection of appropriate risk assessment behaviors and/or coping strategies [11]. Moreover, the distance was not changed by ARS, presumed this behavior does not appear to be driven by general behavioral hyperactivity (an increase on crossings) but rather by a voluntary (brain driven) decision not to visit the periphery zone as non-stressed controls [11]. These findings suggest that stressed zebrafish seem to adopt a coping style that is less suited for successful escape and therefore might not be adaptive (e.g., increases the risk for predator attacks), thus supporting the view that acute stress may impair cognitive abilities. It is also possible that stress could act as a “disorienting” factor causing zebrafish to lose the ability to discriminate between the different parts of the open field [11].

The stimulation of intracellular pathways induced by stress also leads to the free radical generation from intracellular reactive oxygen species, such as superoxide, hydrogen peroxide, and hydroxyl radicals resulting in an imbalance of antioxidant status, disturbances in homeostasis, and oxidative stress [40, 44]. In our study, ARS lowered GSH (NPSH) levels, making the zebrafish brain more susceptible to oxidative damage like increased lipid peroxidation (TBARS). Numerous reports have revealed that restraint stress can affect central nervous system functions by producing neurochemical and hormonal abnormalities associated with oxidative stress in zebrafish [10, 12, 21]. Also, many studies have shown that restraint stress increases lipid peroxidation and increases or decreases antioxidant enzyme activities in different brain regions of rodents depending on the severity and duration of immobilization stress protocol from inducing numerous cellular cascades that lead to increased reactive oxygen species production [40]. In our study, both curcumin preparations were not able to prevent the ARS-induced effects on behavioral and biochemical parameters in zebrafish.

There is controversy in the literature about the anxiolytics effects of curcumin. In rodents subjected to acute immobile stress (120 minutes), both anxiety and hyperactivity were reversed by pre-treatment with curcumin (200 mg/kg/day for 7 days) in the open field test [40]. Also, the pre-treatment with curcumin (20 mg/kg) prevented the hyperlocomotion and anxiogenic state induced by restraint stress (6 hours) in mice [40, 45]. However, in another study, curcumin (20 mg/kg) did not demonstrate any effects in the open field test, and no interaction of curcumin at the benzodiazepine site of the GABAA receptor was observed [46].

Curcumin has recently been classified as both a PAINS (pan-assay interference compounds) and an IMPS (invalid metabolic panaceas) candidate. Curcumin has shown promise in thousands of preclinical studies. However, over 100 clinical trials have failed to find health benefits in humans against several diseases. No form of curcumin, or its closely related analogs, appears to possess the properties required for a good drug candidate (chemical stability, high water solubility, potent and selective target activity, high bioavailability, broad tissue distribution, stable metabolism, and low toxicity). The essential medicinal chemistry of curcumin provides evidence that curcumin is an unstable, reactive, and nonbioavailable compound. Moreover, the available evidence demonstrates that curcumin will ultimately degrade upon release into physiologic media, no matter the delivery mechanism. [47]. Acute curcumin does not seem to have an anti-stress effect on behavior parameters in zebrafish. In our previous study, both curcumin and micronized curcumin were unable to block the behavioral effects of chronic stress (14 days) in zebrafish, even after chronic treatment (7 days), and only antioxidant effects were observed, which were not sufficient to block its behavioral effects [31].

## Conclusions

In this study, we have shown that ARS increased locomotion and decreased thigmotaxis in the open tank test and induced oxidative damage in the zebrafish brain. We suppose that acute stress may impair cognitive abilities acting as a disorienting factor causing a reduction in anxiety-like behaviors such as risk assessment behaviors in zebrafish. Moreover, both curcumin preparations were not able to prevent the effects of ARS on behavioral and biochemical parameters in zebrafish. These results corroborate with previous data showing curcumin (for 7 days) did not seem to have a behavioral effect in zebrafish, even after the micronization process. We speculate that the effects of curcumin on stress-induced behavioral changes might be observed with a longer exposure time or even in a dose range different from that used in these studies.

## References

1. Ulrich-Lai YM, Herman JP (2009) Neural regulation of endocrine and autonomic stress responses. Nat Rev Neurosci 10:397–409. https://doi.org/10.1038/nrn2647

2. Chattarji S, Tomar A, Suvrathan A, et al (2015) Neighborhood matters: divergent patterns of stress-induced plasticity across the brain. Nat Neurosci 18:1364–1375. https://doi.org/10.1038/nn.4115

3. McEwen BS, Bowles NP, Gray JD, et al (2015) Mechanisms of stress in the brain. Nat Neurosci 18:1353–1363. https://doi.org/10.1038/nn.4086

4. McEwen BS (2007) Physiology and neurobiology of stress and adaptation: central role of the brain. Physiol Rev 87:873–904. https://doi.org/10.1152/physrev.00041.2006

5. Joëls M, Baram TZ (2009) The neuro-symphony of stress. Nat Rev Neurosci 10:459–466. https://doi.org/10.1038/nrn2632

6. Miller AH, Raison CL (2016) The role of inflammation in depression: from evolutionary imperative to modern treatment target. Nat Rev Immunol 16:22–34. https://doi.org/10.1038/nri.2015.5

7. Ramaholimihaso T, Bouazzaoui F, Kaladjian A (2020) Curcumin in Depression: Potential Mechanisms of Action and Current Evidence-A Narrative Review. Front Psychiatry 11:572533. https://doi.org/10.3389/fpsyt.2020.572533

8. Alsop D, Vijayan MM (2008) Development of the corticosteroid stress axis and receptor expression in zebrafish. Am J Physiol-Regul Integr Comp Physiol 294:R711–R719. https://doi.org/10.1152/ajpregu.00671.2007

9. Abreu MS de, Koakoski G, Ferreira D, et al (2014) Diazepam and Fluoxetine Decrease the Stress Response in Zebrafish. PLoS ONE 9:e103232. https://doi.org/10.1371/journal.pone.0103232

10. Bertelli PR, Mocelin R, Marcon M, et al (2021) Anti-stress effects of the glucagon-like peptide-1 receptor agonist liraglutide in zebrafish. Prog Neuropsychopharmacol Biol Psychiatry 111:110388. https://doi.org/10.1016/j.pnpbp.2021.110388

11. Champagne DL, Hoefnagels CCM, de Kloet RE, Richardson MK (2010) Translating rodent behavioral repertoire to zebrafish (Danio rerio): Relevance for stress research. Behav Brain Res 214:332–342. https://doi.org/10.1016/j.bbr.2010.06.001

12. Dal Santo G, Conterato GMM, Barcellos LJG, et al (2014) Acute restraint stress induces an imbalance in the oxidative status of the zebrafish brain. Neurosci Lett 558:103–108. https://doi.org/10.1016/j.neulet.2013.11.011

13. Egan RJ, Bergner CL, Hart PC, et al (2009) Understanding behavioral and physiological phenotypes of stress and anxiety in zebrafish. Behav Brain Res 205:38–44. https://doi.org/10.1016/j.bbr.2009.06.022

14. Fontana BD, Cleal M, Gibbon AJ, et al (2021) The effects of two stressors on working memory and cognitive flexibility in zebrafish (Danio rerio): The protective role of D1/D5 agonist on stress responses. Neuropharmacology 196:108681. https://doi.org/10.1016/j.neuropharm.2021.108681

15. Ghisleni G, Capiotti KM, Da Silva RS, et al (2012) The role of CRH in behavioral responses to acute restraint stress in zebrafish. Prog Neuropsychopharmacol Biol Psychiatry 36:176–182. https://doi.org/10.1016/j.pnpbp.2011.08.016

16. Giacomini ACVV, Abreu MS, Giacomini LV, et al (2016) Fluoxetine and diazepam acutely modulate stress induced-behavior. Behav Brain Res 296:301–310. https://doi.org/10.1016/j.bbr.2015.09.027

17. Idalencio R, Kalichak F, Rosa JGS, et al (2015) Waterborne Risperidone Decreases Stress Response in Zebrafish. PLOS ONE 10:e0140800. https://doi.org/10.1371/journal.pone.0140800

18. Mocelin R, Herrmann AP, Marcon M, et al (2015) N-acetylcysteine prevents stress-induced anxiety behavior in zebrafish. Pharmacol Biochem Behav 139:121–126. https://doi.org/10.1016/j.pbb.2015.08.006

19. Pancotto L, Mocelin R, Marcon M, et al (2018) Anxiolytic and anti-stress effects of acute administration of acetyl-L-carnitine in zebrafish. PeerJ 6:e5309. https://doi.org/10.7717/peerj.5309

20. Piato AL, Rosemberg DB, Capiotti KM, et al (2011) Acute restraint stress in zebrafish: behavioral parameters and purinergic signaling. Neurochem Res 36:1876–1886. https://doi.org/10.1007/s11064-011-0509-z

21. Reis CG, Mocelin R, Benvenutti R, et al (2020) Effects of N-acetylcysteine amide on anxiety and stress behavior in zebrafish. Naunyn Schmiedebergs Arch Pharmacol 393:591–601. https://doi.org/10.1007/s00210-019-01762-8

22. da Silva Marques JG, Antunes FTT, da Silva Brum LF, et al (2021) Adaptogenic effects of curcumin on depression induced by moderate and unpredictable chronic stress in mice. Behav Brain Res 399:113002. https://doi.org/10.1016/j.bbr.2020.113002

23. Khodadadegan MA, Azami S, Guest PC, et al (2021) Effects of Curcumin on Depression and Anxiety: A Narrative Review of the Recent Clinical Data. Adv Exp Med Biol 1291:283–294. https://doi.org/10.1007/978-3-030-56153-6_17

24. Lopresti AL, Maes M, Maker GL, et al (2014) Curcumin for the treatment of major depression: A randomised, double-blind, placebo controlled study. J Affect Disord 167:368–375. https://doi.org/10.1016/j.jad.2014.06.001

25. Matias JN, Achete G, Campanari GSDS, et al (2021) A systematic review of the antidepressant effects of curcumin: Beyond monoamines theory. Aust N Z J Psychiatry 55:451–462. https://doi.org/10.1177/0004867421998795

26. Mohammad Abu-Taweel G, Al-Fifi Z (2021) Protective effects of curcumin towards anxiety and depression-like behaviors induced mercury chloride. Saudi J Biol Sci 28:125–134. https://doi.org/10.1016/j.sjbs.2020.09.011

27. Yang K-Y, Lin L-C, Tseng T-Y, et al (2007) Oral bioavailability of curcumin in rat and the herbal analysis from Curcuma longa by LC–MS/MS. J Chromatogr B 853:183–189. https://doi.org/10.1016/j.jchromb.2007.03.010

28. Anand P, Kunnumakkara AB, Newman RA, Aggarwal BB (2007) Bioavailability of Curcumin: Problems and Promises. Mol Pharm 4:807–818. https://doi.org/10.1021/mp700113r

29. Almeida ER, Lima-Rezende CA, Schneider SE, et al (2021) Micronized Resveratrol Shows Anticonvulsant Properties in Pentylenetetrazole-Induced Seizure Model in Adult Zebrafish. Neurochem Res 46:241–251. https://doi.org/10.1007/s11064-020-03158-0

30. Decui L, Garbinato CLL, Schneider SE, et al (2020) Micronized resveratrol shows promising effects in a seizure model in zebrafish and signalizes an important advance in epilepsy treatment. Epilepsy Res 159:106243. https://doi.org/10.1016/j.eplepsyres.2019.106243

31. Sachett A, Gallas-Lopes M, Benvenutti R, et al (2021) Curcumin micronization by supercritical fluid: in vitro and in vivo biological relevance. bioRxiv 2021.07.08.451641. https://doi.org/10.1101/2021.07.08.451641

32. Sert NP du, Hurst V, Ahluwalia A, et al (2020) The ARRIVE guidelines 2.0: Updated guidelines for reporting animal research. Br J Pharmacol 177:3617–3624. https://doi.org/10.1111/bph.15193

33. Leary S, Pharmaceuticals F, Underwood W, et al (2020) AVMA Guidelines for the Euthanasia of Animals: 2020 Edition. 121

34. Benvenutti R, Gallas-Lopes M, Sachett A, et al (2020) How do zebrafish respond to MK-801 and amphetamine? Relevance for assessing schizophrenia-relevant endophenotypes in alternative model organisms. bioRxiv 2020.08.03.234567. https://doi.org/10.1101/2020.08.03.234567

35. Sachett A (2020) How to prepare zebrafish brain tissue samples for biochemical assays. https://doi.org/10.17504/protocols.io.bjkdkks6

36. Bradford MM (1976) A rapid and sensitive method for the quantitation of microgram quantities of protein utilizing the principle of protein-dye binding. Anal Biochem 72:248–254. https://doi.org/10.1006/abio.1976.9999

37. Sachett A (2020) Protein quantification protocol optimized for zebrafish brain tissue (Bradford method). https://doi.org/10.17504/protocols.io.bjnfkmbn

38. Sachett A, Gallas-Lopes M, Conterato GMM, et al (2021) Quantification of nonprotein sulfhydryl groups (NPSH) optimized for zebrafish brain tissue. In: protocols.io. https://www.protocols.io/view/quantification-of-nonprotein-sulfhydryl-groups-nps-bx8tprwn. Accessed 11 Oct 2021

39. Sachett A (2020) Quantification of thiobarbituric acid reactive species (TBARS) optimized for zebrafish brain tissue. https://doi.org/10.17504/protocols.io.bjp8kmrw

40. Haider S, Naqvi F, Batool Z, et al (2015) Pretreatment with curcumin attenuates anxiety while strengthens memory performance after one short stress experience in male rats. Brain Res Bull 115:1–8. https://doi.org/10.1016/j.brainresbull.2015.04.001

41. Johnson A, Hamilton TJ (2017) Modafinil decreases anxiety-like behaviour in zebrafish. PeerJ 5:e2994. https://doi.org/10.7717/peerj.2994

42. Stewart A, Gaikwad S, Kyzar E, et al (2012) Modeling anxiety using adult zebrafish: A conceptual review. Neuropharmacology 62:135–143. https://doi.org/10.1016/j.neuropharm.2011.07.037

43. McEwen BS, Wingfield JC (2003) The concept of allostasis in biology and biomedicine. Horm Behav 43:2–15. https://doi.org/10.1016/s0018-506x(02)00024-7

44. Uttara B, Singh AV, Zamboni P, Mahajan RT (2009) Oxidative Stress and Neurodegenerative Diseases: A Review of Upstream and Downstream Antioxidant Therapeutic Options. Curr Neuropharmacol 7:65–74. https://doi.org/10.2174/157015909787602823

45. Gilhotra N, Dhingra D (2010) GABAergic and nitriergic modulation by curcumin for its antianxiety-like activity in mice. Brain Res 1352:167–175. https://doi.org/10.1016/j.brainres.2010.07.007

46. Ceremuga TE, Helmrick K, Kufahl Z, et al (2017) Investigation of the Anxiolytic and Antidepressant Effects of Curcumin, a Compound From Turmeric (Curcuma longa), in the Adult Male Sprague-Dawley Rat. Holist Nurs Pract 31:193–203. https://doi.org/10.1097/HNP.0000000000000208

47. Nelson KM, Dahlin JL, Bisson J, et al (2017) The Essential Medicinal Chemistry of Curcumin. J Med Chem 60:1620–1637. https://doi.org/10.1021/acs.jmedchem.6b00975

